# Tackling the Complexity of Spatial Transcriptomics Data Interpretation with Large Language Models

**DOI:** 10.1101/2024.11.28.625773

**Authors:** Taushif Khan, Colleen M. Farley, John J. Wilson, Chih-Hao Chang, Damien Chaussabel

## Abstract

Spatial transcriptomics offers unprecedented insights into the complex cellular landscapes of tissues, particularly in cancer research where understanding the tumor microenvironment is crucial. However, interpreting the vast and intricate data generated by this technology remains a significant challenge. This study explores the potential of Large Language Models (LLMs) to assist in the analysis and interpretation of spatial transcriptomic data from a murine melanoma tumor model. We first evaluated the performance of multiple LLM models in describing and quantifying spatial gene expression patterns. Our benchmarking revealed that spatial transcriptomics data interpretation proved challenging for most models, with only a few demonstrating sufficient capability for this complex task. Using Claude 3.5 Sonnet, which showed the highest accuracy in spot quantification and pattern recognition, we developed a systematic workflow for analyzing the tumor immune landscape. The model first assisted in identifying and prioritizing panels of M1 and M2 macrophage-associated markers through knowledge-driven scoring. It then demonstrated remarkable ability to integrate spatial expression data with extensive immunological knowledge, providing sophisticated interpretation of local immune organization. When analyzing individual tumor regions, the model identified coordinated immunosuppressive mechanisms including metabolic barriers and disrupted pro-inflammatory signaling cascades, findings that both aligned with and extended current understanding of tumor immunology. This study highlights the potential of LLMs as powerful assistive tools in spatial transcriptomics analysis, capable of combining advanced pattern recognition with extensive knowledge integration to enhance data interpretation. While significant development work remains to make such workflows scalable, our proof of concept demonstrates that LLMs can help accelerate the translation of spatial transcriptomics data into biological insights.

## INTRODUCTION

Spatial transcriptomics has emerged as a transformative technology in the field of molecular biology, offering unprecedented insights into the spatial organization and function of cells within complex tissues. Unlike traditional bulk RNA sequencing methods, which provide an averaged gene expression profile across a tissue sample, spatial transcriptomics preserves the spatial context of gene expression, allowing researchers to map transcriptional activity to specific locations within a tissue section (1). This technology has found wide-ranging applications, from developmental biology to cancer research, providing a high-resolution view of cellular heterogeneity and interactions in their native microenvironment (2). In cancer research, spatial transcriptomics has been particularly valuable in elucidating the complex interactions within the tumor microenvironment, revealing spatial patterns of immune cell infiltration, stromal cell distribution, and tumor cell heterogeneity (3).

However, the interpretation of this complex, multidimensional data remains a significant challenge, requiring sophisticated computational approaches and deep biological knowledge to extract meaningful insights (4). Large Language Models (LLMs) have recently emerged as powerful tools with the potential to address this challenge. These models, trained on extensive textual data, demonstrate remarkable abilities to process and understand complex information across various domains (5,6). In the field of biomedical research, LLMs have shown promise in tasks such as literature mining, hypothesis generation, and data interpretation (7). Recent work has demonstrated the utility of LLMs in candidate gene prioritization and selection, showcasing their ability to leverage extensive biomedical knowledge for focused analytical tasks (8). Further advancements have equipped LLMs with functionalities to interpret visual data, including scientific images and figures (9). This capability is particularly relevant for spatial transcriptomics, where the visual representation of gene expression patterns is crucial. The potential of LLMs to integrate textual knowledge with visual interpretation offers a novel approach to assist researchers in deciphering the complex spatial patterns of gene expression within tissues.

In this study, we explore the application of LLMs to the analysis and interpretation of spatial transcriptomics data from a murine melanoma tumor model. We first assess different LLM architectures for their ability to quantify spatial gene expression patterns, then develop a systematic workflow combining spatial analysis with knowledge-driven marker selection to characterize the immune landscape within tumor regions. Through this proof of concept, we aim to demonstrate how LLMs might assist researchers in extracting mechanistic insights from complex spatial molecular data.

## METHODS

### Murine Melanoma Model Preparation

#### Mice

C57BL/6J mice (The Jackson Laboratory, JAX) were housed in a specific-pathogen-free animal facility at 22°C, between 40-50% humidity, with a daylight cycle from 6:00 am to 6:00 pm. Mice were provided 6% fat food (LabDiet 5K52) and water ad libitum. Age-matched female B6 mice between 8-10 weeks old were used for experiments. Animal experiments were conducted in accordance with procedures approved by the Institutional Animal Care and Use Committee of The Jackson Laboratory (#17070-1).

### Cell Lines

Murine melanoma cells, B16-OVA (provided by Dr. Hildehund Ertl, with permission from Drs. Edith Lord and John Frelinger), were authenticated by whole genome and transcriptome sequencing. B16-OVA cells were transduced with lentivirus expressing mCherry fluorescent protein and overexpressing hexokinase II (HK2, B16-HK2) or empty vector control (B16-Mock). Transduced cells were sort purified for mCherry expression and, for B16-HK2 cells, high glycolytic activity (determined by fluorescent 2- NBDG uptake). Cells were grown in RPMI-1640 medium supplemented with penicillin, streptomycin, and 10% FBS. HK2-mCherry cells were routinely screened for HK2 overexpression using anti-Hexokinase II antibody (Abcam, ab209847) by western blot.

### Tumor Model and Treatment

5 x 10^5^ B16-OVA derivative cells were subcutaneously injected into the right flank of B6 mice. Anti-PD-1 antibody (200 μg/mouse, Leinco Technologies Inc., P362) or isotype control antibody was intraperitoneally injected every 3 days, beginning 5 days post-tumor inoculation, until harvest on day 15. Tumors were dissected, embedded in OCT medium, and stored at - 80°C until sectioning.

### Spatial Transcriptomics Sample Preparation

#### Visium Slide Preparation

OCT-embedded tumor blocks were cryosectioned at 10 μm thickness. Sections were placed on Visium Spatial slides (10x Genomics, 2000233) within capture areas. RNA integrity was assessed using the Agilent RNA 6000 Nano Kit on the 2100 Bioanalyzer system, with all samples achieving a RNA Integrity Number (RIN) ≥ 9.0.

#### Data Processing and Visualization

Raw sequencing data were processed using the Space Ranger pipeline (10x Genomics) to generate gene expression matrices and spatial feature plots. The Seurat R package was used for downstream analysis and visualization of spatial gene expression patterns.

### LLM evaluation and prompt optimization

#### Model selection and evaluation

Initial evaluations were conducted in March-April 2024 using models from three major AI companies (Anthropic, OpenAI, and Google) that offered image analysis capabilities through their chat interfaces at that time. Models were accessed through their respective web interfaces. Subsequent optimization and comparative analyses were performed in October 2024, including a wider model offering.

#### Prompt design and iteration

We developed and evaluated three different prompts for quantitative analysis of spatial transcriptomics data: *Prompt A (Initial comprehensive prompt – adopted Q2 2024):* For each grid square (A1 to E5), models were instructed to: 1) estimate the total number of spots; 2) count spots with high expression (yellow), medium expression (green), and low/no expression (purple); 3) present the results in a summary table with columns for Grid Square ID, High Expression (Yellow), Medium Expression (Green), Low/No Expression (Purple), and Total Spots. Models were also asked to calculate the overall percentage distribution of expression levels across the entire tumor section and estimate the percentage area covered by each expression level in each grid square.

*Prompt B (Optimized structured stepwise prompt – adopted Q4 2024):* "Analyze the spatial distribution of markers across the tumor section shown in the attached image. Follow these steps for each grid square from A1 to E5:

1. Estimate the total number of spots in the square
2. Count the number of yellow spots (high expression)
3. Count the number of green spots (medium expression)
4. Count the number of purple spots (low to no expression)
5. Verify that the sum of yellow, green, and purple spots equals the total from step 1.

If not, recheck your counts

For each grid square, record:

- Square ID
- Total spot count
- Yellow spot count
- Green spot count
- Purple spot count

Present your findings in a table with these five columns. Include a note on any squares where spot counts were particularly challenging to estimate or where you noticed interesting patterns. After completing the table, sum up the total spots and the counts for each color across all squares. Use these sums to calculate the percentage of each color, rounding to the nearest whole percent. Finally, identify the three squares with the highest total spot counts and the three with the lowest, explaining any patterns you observe in their locations or color distributions."

#### LLM Approach for M1 and M2 Marker Identification

Following the benchmarking of LLMs for spatial transcriptomics data interpretation, we employed a multi-step LLM approach to identify and prioritize markers associated with M1 and M2 macrophage phenotypes in the tumor microenvironment.

#### Cross-LLM Marker Screen

We initiated a broad screening process using multiple LLMs to identify potential M1 and M2 macrophage markers. The LLMs employed in this screen included Mistral AI, GPT-4o, Claude 3.5, Consensus GPT, Gemini Ultra, and Cohere AI. Each model was presented with two successive prompts for both M1 and M2 macrophages. For M1 macrophages, the first prompt was: "Provide a list of markers for murine (mouse) M1 macrophages in the tumor microenvironment. Output a table with the first column indicating the official gene symbol, the second column a brief justification."

The second prompt requested: "Provide in addition a list of less canonical murine M1 macrophage markers." Analogous prompts were used for M2 macrophages, substituting "M1" with "M2". The outputs from each LLM were consolidated into comprehensive lists of potential M1 and M2 markers for further evaluation.

#### LLM Scoring of Macrophage Markers

Following the initial identification, we employed GPT-4o and Claude 3.5 to score and prioritize the identified markers. For each marker, both LLMs were presented with a detailed prompt that included the following tasks:

1. Provide the gene’s official name
2. Provide a brief summary of the gene’s function
3. Score each of the following statements from 0 to 10, with 0 indicating no evidence and 10 indicating conclusive evidence: a. The gene’s expression is a marker of M1 macrophages b. The gene’s product stimulates immune responses in the tumor microenvironment c. The gene’s expression is a marker of M2 macrophages d. The gene’s product suppresses immune responses in the tumor microenvironment

The LLMs were instructed to use a specific scoring criteria, ranging from 0 (no evidence found) to 10 (strong evidence), and to provide an evaluative comment and up to three supporting references from peer-reviewed literature for scores of 4 or above.

### Marker Prioritization and Selection

We developed a ranking system based on the averaged scores provided by GPT-4o and Claude 3.5. For each marker, we calculated:

1. An M1 score by averaging scores for statements a and b (strength of M1 association)
2. An M2 score by averaging scores for statements c and d (strength of M2 association)
3. Specificity ratios:

a. For M1 markers: M1 score / M2 score
b. For M2 markers: M2 score / M1 score

Markers were then ranked separately based on:

1. Their association strength (M1 or M2 score)
2. Their specificity ratio

The final rank for each marker was determined by summing these two rank values, with lower sums indicating higher overall ranks. This process was carried out separately for M1 and M2 markers, aiming to identify 15 M1 markers and 14 M2 markers.

We excluded markers that showed low specificity or contradictory evidence. Specifically, Arg1 was excluded from the M1 marker list, and Cdh1 was excluded from the M2 marker list based on their scoring averages and specificity ratios. Additionally, we eliminated redundancies (e.g., when both IL12a and IL12 appeared among top-ranked genes) and resolved cases of ambiguous nomenclature (e.g., MHC-II).

This multi-step LLM approach allowed us to leverage the broad knowledge base of multiple models for initial identification, followed by more focused scoring and prioritization using the top-performing models. The result was a refined list of 15 M1 and 14 M2 macrophage markers, ranked by their likelihood of being reliable indicators of macrophage phenotypes in the tumor microenvironment.

#### Claude 3.5 Interpretation and Immune Status Scoring of Spatial Transcriptomic Data

Following the identification and prioritization of M1 and M2 macrophage markers, we evaluated the ability of Claude 3.5 Sonnet to interpret spatial transcriptomic data and generate an immune status score for specific grid sections of the tumor.

### Spatial Transcriptomic Data Interpretation

We focused on grid position A2 of the murine melanoma tumor section for this analysis. Claude 3.5 Sonnet was presented with expression data for 30 markers (15 M1 and 15 M2) sequentially. The LLM was first provided with the following context: "We have used Visium technology to generate spatial transcriptomic data from a murine melanoma tumor section. Our aim is to achieve an intricate mapping of the tumor microenvironment, focusing particularly on the interplay between macrophage-mediated immune suppression and activation. The grid in the attached image represents the tumor section, delineated by coordinates A through E and 1 through 5."

Next, for each marker, two images were provided as input: the expression pattern for grid section A2 and the corresponding marker saturation scale. Claude 3.5 Sonnet was then prompted with: "Focusing on A2 and the [first/next] marker associated with macrophage-mediated [immune suppression/immune stimulation], here is the expression of [marker name]. Generate a table that will be appended as we go, starting with this first marker, and columns indicating: the total number of spots; the number of yellow spots; the number of green spots; the number of purple spots; the spatial distribution of yellow, green and purple spots, the saturation scale for this marker."

### Immune Status Scoring

After analyzing all markers, we tasked Claude 3.5 Sonnet with assigning an immune status score to the A2 grid section. The LLM was instructed to use a scale ranging from -10 to +10, with -10 denoting maximal immunosuppression and +10 indicating maximal immune activation. The prompt was: "On a scale ranging from - 10 to +10, with -10 denoting maximal immunosuppression and +10 indicating maximal immune activation, please assign a score to the A2 grid section of the murine melanoma tumor. This score should reflect the cumulative immunological state as determined by the spatial expression of the markers analyzed."

### Knowledge-Driven Weighting Approach

Finally, we employed a knowledge-driven weighting approach to further refine the immune status score. Claude 3.5 Sonnet was instructed to weight the functional significance of each marker in the context of the tumor microenvironment and cancer immunotherapy. Markers were categorized as highly significant (weight: 3), moderately significant (weight: 2), or less significant (weight: 1). Absent markers of high significance were assigned a weight of -3 for immunostimulatory markers and +3 for immunosuppressive markers. The LLM was prompted to calculate a weighted score considering these functional weights alongside the expression levels and spatial distributions of the markers.

#### Comprehensive Characterization of Immunological Phenotype

Following spot quantification, spatial pattern analysis, and functionally-weighted scoring, we tasked Claude 3.5 Sonnet with providing an integrated characterization of the immunological phenotype for grid section A2. The LLM was first prompted to synthesize all previous analyses: "Based on the quantitative assessment of spatial distribution patterns, functional significance of markers, and pathway relationships previously analyzed, please provide a comprehensive characterization of the immunological phenotype for grid section A2. Consider both the abundance and spatial organization of all 30 markers, their functional hierarchies, and their coordinated effects on the immune microenvironment." The model was then asked to contextualize these observations within the current state of knowledge and to provide a narrative summary supported by literature references.

## RESULTS

### Spatial Transcriptomics Reveals Complex Organization of the Murine Melanoma Microenvironment

To establish a framework for evaluating LLM capabilities in spatial transcriptomics analysis, we first generated high-quality spatial gene expression data from murine B16- OVA melanoma tumors using the 10x Genomics Visium platform. The tumor section exhibited excellent RNA quality (RIN ≥ 9.0) and contained approximately 2,000 spatially-resolved spots distributed across a grid-defined area (A1-E5) (Fig. 1A). Each spot, representing a discrete spatial unit of approximately 55 μm in diameter, captured transcripts from an estimated 10-20 cells. Manual quantification of expression patterns across the tumor section revealed distinct spatial organization, with clear gradients and localized regions of differential expression (Fig. 1A, right panel). The expression data showed high technical quality, with consistent spot detection across the tissue and clear distinction between tissue-containing and empty spots. This spatial transcriptomics dataset thus provided a robust foundation for evaluating the capability of LLMs to interpret complex spatial gene expression patterns in the tumor microenvironment.

**Figure 1:**
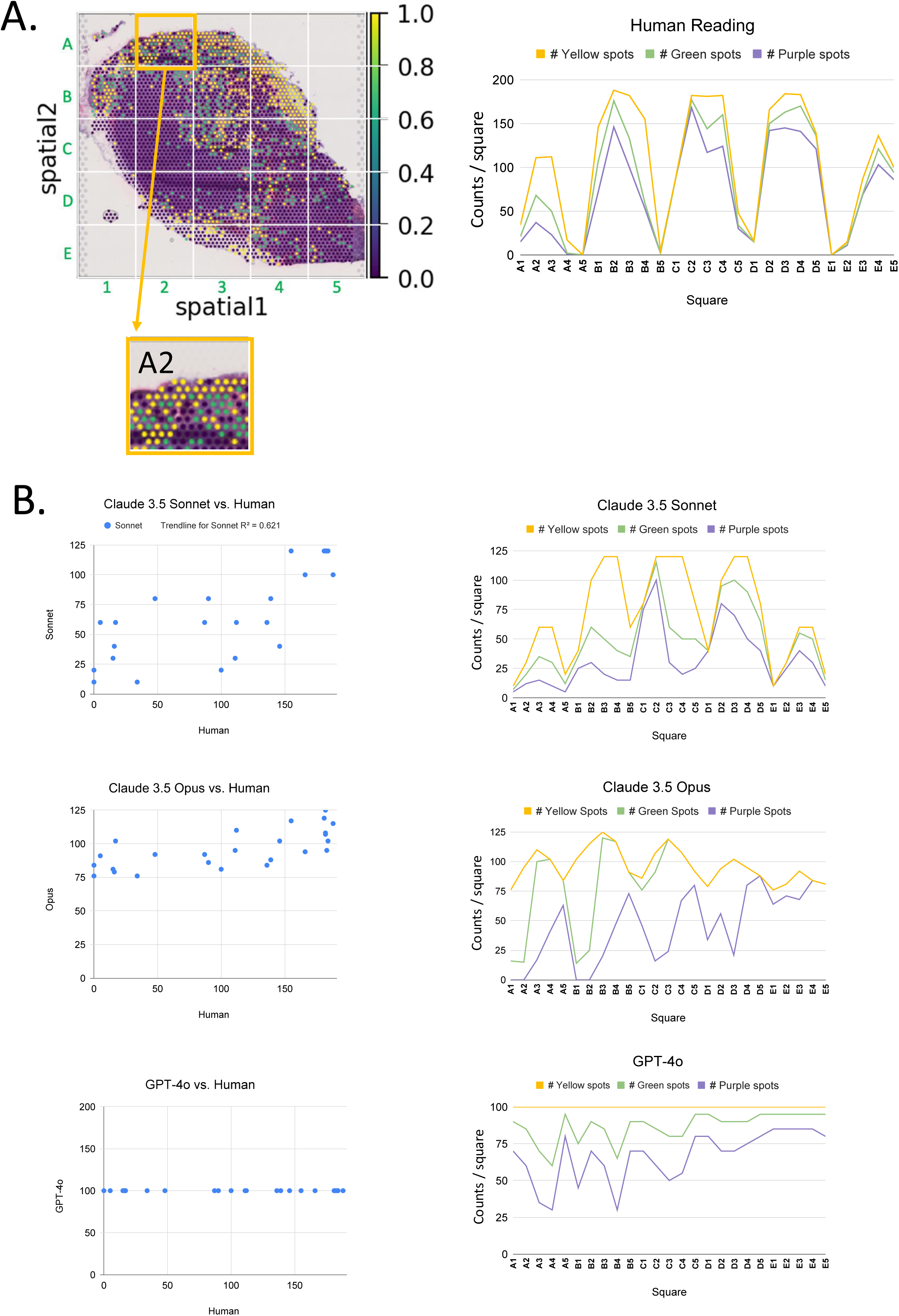
Benchmarking LLM performance in spatial transcriptomics spot quantification. **A.** Representative spatial transcriptomics data from a murine melanoma tumor section showing marker expression patterns. Left: Grid overlay (A1-E5) on tumor section with color scale indicating expression levels from 0.0 (purple) to 1.0 (yellow). Grid section A2 is highlighted (yellow box) and shown at higher magnification below. Right: Human expert quantification of spots across grid sections, showing distribution of high (yellow), medium (green), and low (purple) expression spots. **B.** Performance comparison of different LLM models in spot quantification. Left panels: Correlation plots showing the relationship between LLM and human expert spot counts for Claude 3.5 Sonnet (R^2^ = 0.621), Claude 3.5 Opus, and GPT-4o. Right panels: Corresponding spot distribution patterns generated by each model across grid sections, displaying counts of high (yellow), medium (green), and low (purple) expression spots.

### Preliminary Benchmarking of LLM Performance in Spatial Transcriptomics Data Interpretation

The ability of LLMs to accurately interpret scientific data visualizations, particularly complex spatial expression patterns, represents an untested frontier in artificial intelligence. To establish a quantitative framework for evaluating LLM performance in this domain, we compared spot-counting accuracy between three leading LLM models - Claude 3.5 Sonnet, Claude 3.5 Opus, and GPT-4o - against manual expert quantification as a gold standard (see Methods). Claude 3.5 Sonnet demonstrated the strongest correlation with expert counting (R^2^ = 0.621), accurately capturing both the general distribution patterns and local variations in expression levels (Fig. 1B). In contrast, Claude 3.5 Opus showed weaker correlation and tendency to overestimate spot counts, while GPT-4o exhibited minimal variation in its estimates across different grid positions, suggesting limited ability to discern spatial patterns. These initial results identified Claude 3.5 Sonnet as the most suitable model for subsequent detailed analysis of spatial transcriptomics data.

### Comprehensive Evaluation of LLM Capabilities in Spatial Pattern Recognition

Given these promising results obtained at the end of Q2 of 2024, we revisited this project and conducted a more extensive evaluation of LLM performance in spatial transcriptomics interpretation in Q4 of 2024. Using six independent replicates, we assessed correlation coefficients (R^2^) between LLM counts and expert quantification across a broader range of available models (see Methods). The updated Claude 3.5 Sonnet (late October 2024 release) achieved the highest median R^2^ of 0.65, showing improved performance over its previous version (early October 2024 - **Fig. 2A**). Other models demonstrated varying capabilities, with Gemini Ultra 1.0 and Claude 3.5 Opus showing moderate performance (median R^2^ ∼0.45), while GPT-4o and Mistral exhibited lower correlation values (R^2^ < 0.3). Detailed analysis of spot distribution patterns revealed that Claude 3.5 Sonnet not only maintained high correlation with manual counting (R^2^ = 0.6051) but also accurately preserved the relative proportions of high, medium, and low expression spots across grid positions (**Fig. 2B**). In contrast, while Gemini Ultra showed reasonable pattern recognition (R^2^ = 0.4327), it tended to underestimate total spot counts. GPT-4o (R^2^ = 0.1294) demonstrated limited ability to distinguish between expression levels, producing more uniform distributions that poorly reflected the actual spatial heterogeneity of the tumor microenvironment. These results not only validated our initial findings but also revealed rapid progress in LLM capabilities for scientific data interpretation, with newer model iterations showing marked improvements in spatial pattern recognition accuracy.

**Figure 2:**
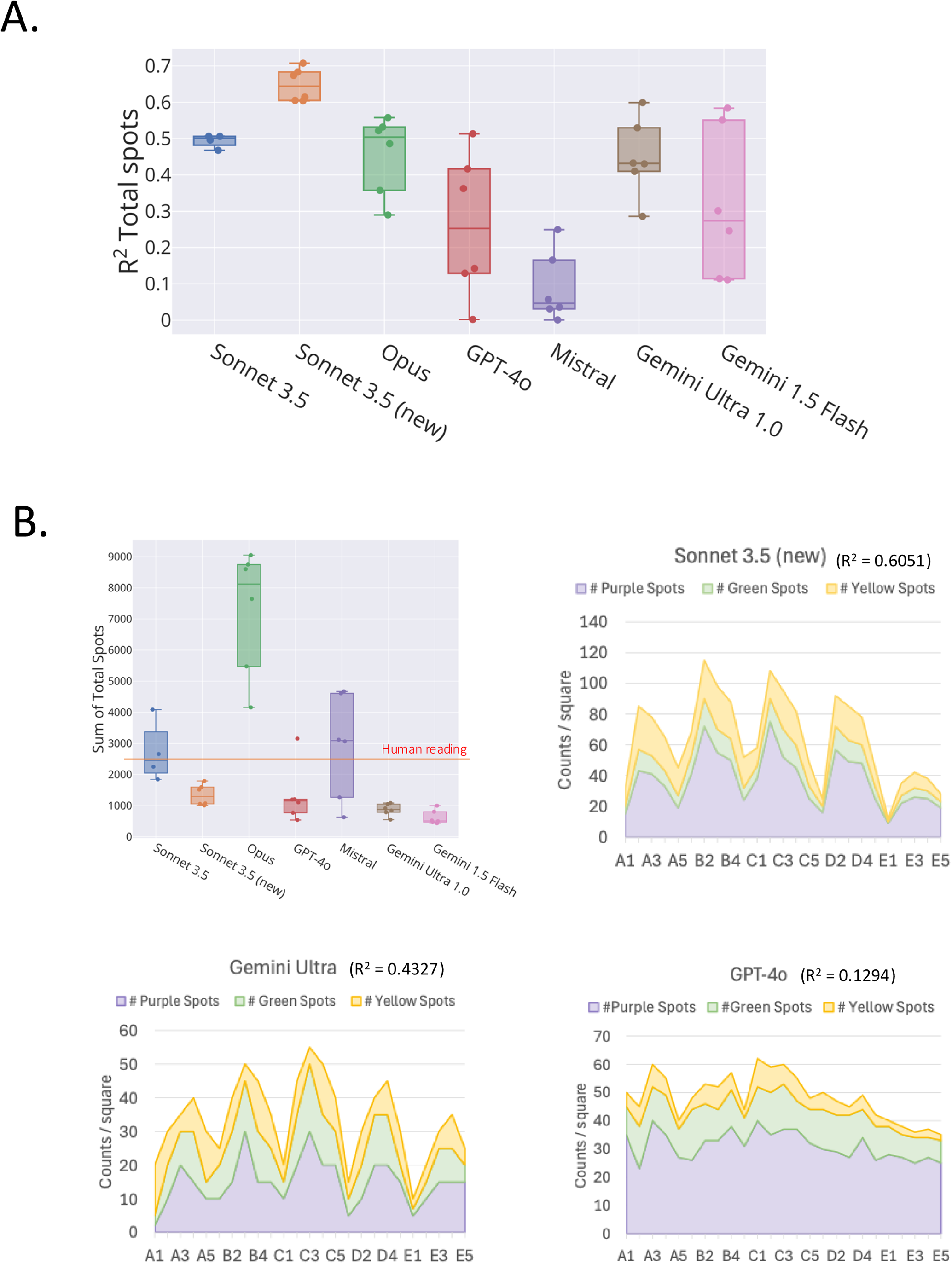
Extended validation of LLM performance across multiple models and replicates. **A.** Box plot showing correlation coefficients (R^2^) between LLM-generated and expert spot counts across six independent replicates. Models tested include Sonnet 3.5, Sonnet 3.5 (new - October 2024 release), Claude 3.5 Opus, GPT-4o, Mistral, Gemini Ultra 1.0, and Gemini 1.5 Flash. Individual data points represent correlation values from separate analysis runs. **B.** Detailed analysis of spot counting accuracy across LLM models. Left: Total spot counts generated by each model compared to human expert quantification (red line). Right panels: Representative spot distribution patterns for the three leading models (Sonnet 3.5 new, Gemini Ultra, GPT-4o) showing counts of high (yellow), medium (green), and low (purple) expression spots across grid sections. Corresponding R^2^ values are indicated for each model.

#### Optimization of LLM Performance Through Multi-step Analysis

While the initial benchmarking established Claude 3.5 Sonnet’s capability in analyzing spatial transcriptomics data, we hypothesized that breaking down the analysis into smaller spatial units might improve accuracy. We therefore implemented a square-by- square analysis strategy, prompting the LLM to examine each grid position individually rather than analyzing the entire tumor section at once (see Methods). This approach substantially improved performance, with correlation coefficients between Sonnet and human quantification increasing from R^2^ = 0.60 to R^2^ = 0.86 (Fig. 3). Notably, the square- by-square analysis also yielded more consistent spot distribution patterns across expression levels, closely matching the relative proportions of high, medium, and low expression spots observed in manual counting. These results demonstrate that appropriate prompting strategies can significantly enhance LLM performance in complex spatial data analysis tasks.

**Figure 3:**
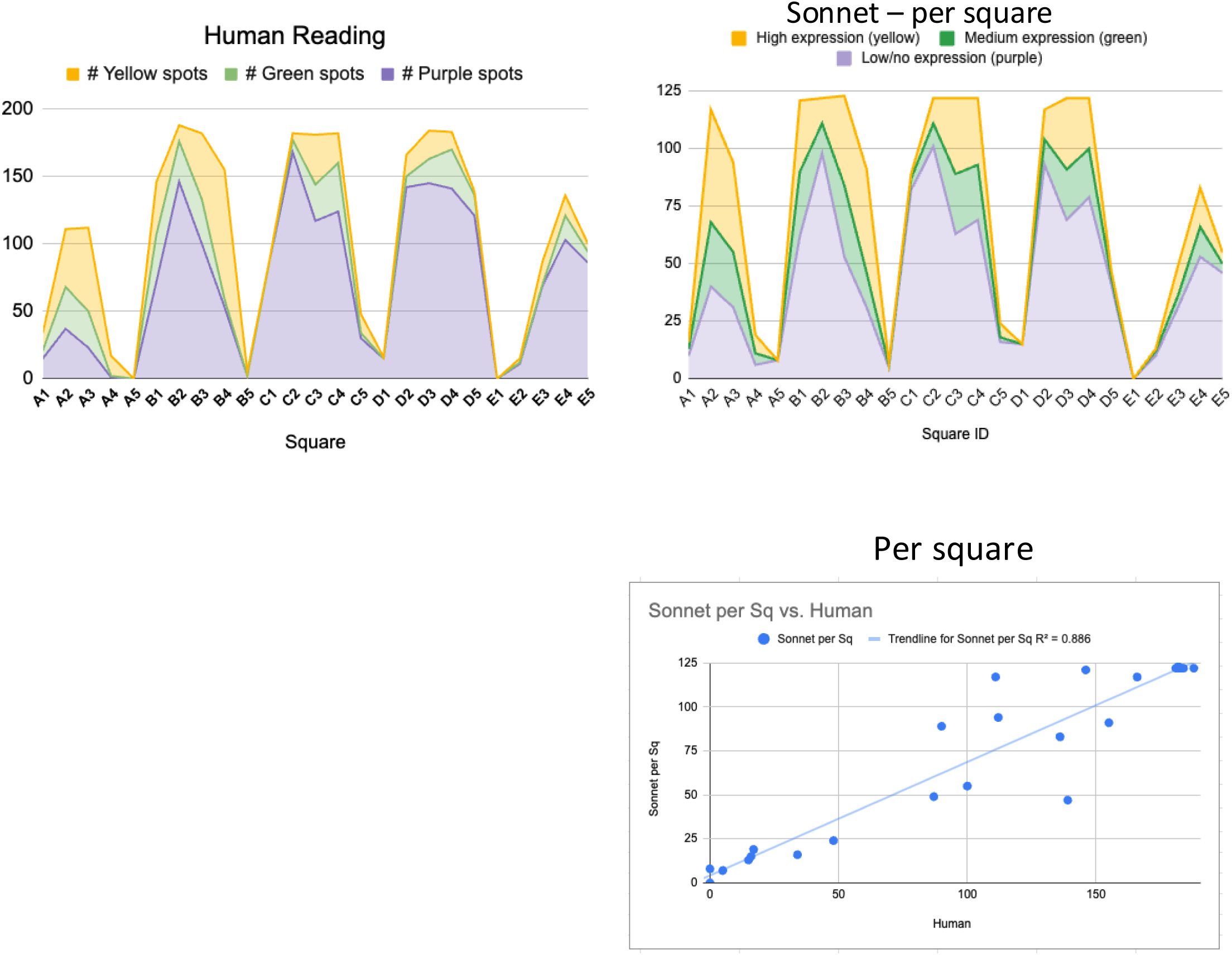
Optimization of LLM analysis through square-by-square assessment of marker expression patterns. Top panels: Representative spatial distribution plots comparing human expert (left) and Claude 3.5 Sonnet (right) spot quantification across grid section A2. Data were obtained through stepwise analysis, where human expert and LLM independently analyzed each grid square individually. Both plots display counts per square for high (yellow), medium (green), and low (purple) expression spots. Note the consistent pattern recognition between human and LLM analysis. Bottom panel: Correlation analysis between Sonnet and human expert spot counts obtained through square-by-square prompting strategy. Scatter plot shows strong correlation (R^2^ = 0.865) between LLM-generated and human spot counts per square, demonstrating improved accuracy achieved through this methodical analytical approach compared to whole-section analysis.

#### LLM-driven Identification and Prioritization of M1 and M2 Macrophage Markers

To characterize the immune microenvironment of the murine melanoma tumor sections, we first sought to establish panels of markers that could reliably distinguish M1 and M2 macrophage populations. We implemented a systematic LLM-based approach combining broad candidate identification with targeted scoring and ranking (see Methods). Initial screening across six different LLMs yielded a comprehensive pool of candidate markers, which were then scored by GPT-4 and Claude 3.5 based on their association with macrophage phenotypes and immunomodulatory functions. Among M1-associated markers, IL12, Cxcl10, and Cxcl9 emerged as the top candidates, with averaged LLM scores of 8.3, 8.5, and 7.5 respectively, and high M1/M2 specificity ratios (6.4, 5.7, and 5.9). These chemokines and cytokines are characteristic of pro-inflammatory, anti-tumoral macrophage responses. For M2-associated markers, IL10, Arg1, and CD163 ranked highest, with notably high LLM scores (9.5, 9.3, and 9.3) and M2/M1 specificity ratios (12.7, 7.0, and 7.7). This set includes canonical M2 markers involved in immunosuppression and tissue remodeling. The relative rankings incorporated both the strength of association with a given phenotype and the specificity of this association, allowing us to exclude markers showing ambiguous or contradictory evidence.

#### Comprehensive Analysis of Immune Marker Expression Patterns in Individual Tumor Regions

To achieve detailed immunological mapping of the tumor microenvironment, we tasked Claude 3.5 Sonnet with analyzing the spatial distribution of the selected M1 and M2 markers within individual grid sections. Using grid section A2 as proof of concept, we sequentially presented the LLM with expression data for each marker, accompanied by their respective saturation scales. The model systematically quantified spot numbers and characterized spatial distributions, generating comprehensive tables for both M2 (**Table 4** & **Figure 4**) and M1 (**Table 5** & **Figure 5**) markers. Claude 3.5 generated the following summary: “Among M2 markers, Hmox1 showed the highest expression (approximately 30 yellow spots on a 0.0-1.5 scale), followed by strong expression of Arg1, Tgfb1, Stab1, and Mrc1 (14-18 yellow spots each). Several M2 markers showed notably low or absent expression, including Alox15 and Cd301. For M1 markers, Stat1 and Cxcl10 demonstrated the strongest expression (approximately 25-30 yellow spots on a 0.0-2.0 scale), while multiple markers including IL12, Tbx21, and Cxcl11 were undetectable. Notably, the LLM’s spot quantification showed high internal consistency and careful consideration of varying saturation scales across markers. The spatial distribution analysis revealed distinct patterns of marker expression across the A2 grid section. High-expressing M2 markers showed a notable concentration in the upper portion of the section, with Arg1 displaying a clear top-to-bottom gradient (18 yellow, 12 green spots). This pattern was partially mirrored by Stab1 and Mrc1, while Tgfb1 showed a more uniform distribution (15 yellow, 12 green spots). Among M1 markers, Stat1 exhibited the most uniform distribution across the section (30 yellow, 10 green spots), while Cxcl10 showed stronger expression in the upper region (25 yellow, 12 green spots). Most low-expressing markers of both types displayed scattered patterns without clear spatial organization, exemplified by IL10 (4 yellow spots) and Tnf (4 green spots only).”

**Figure 4:**
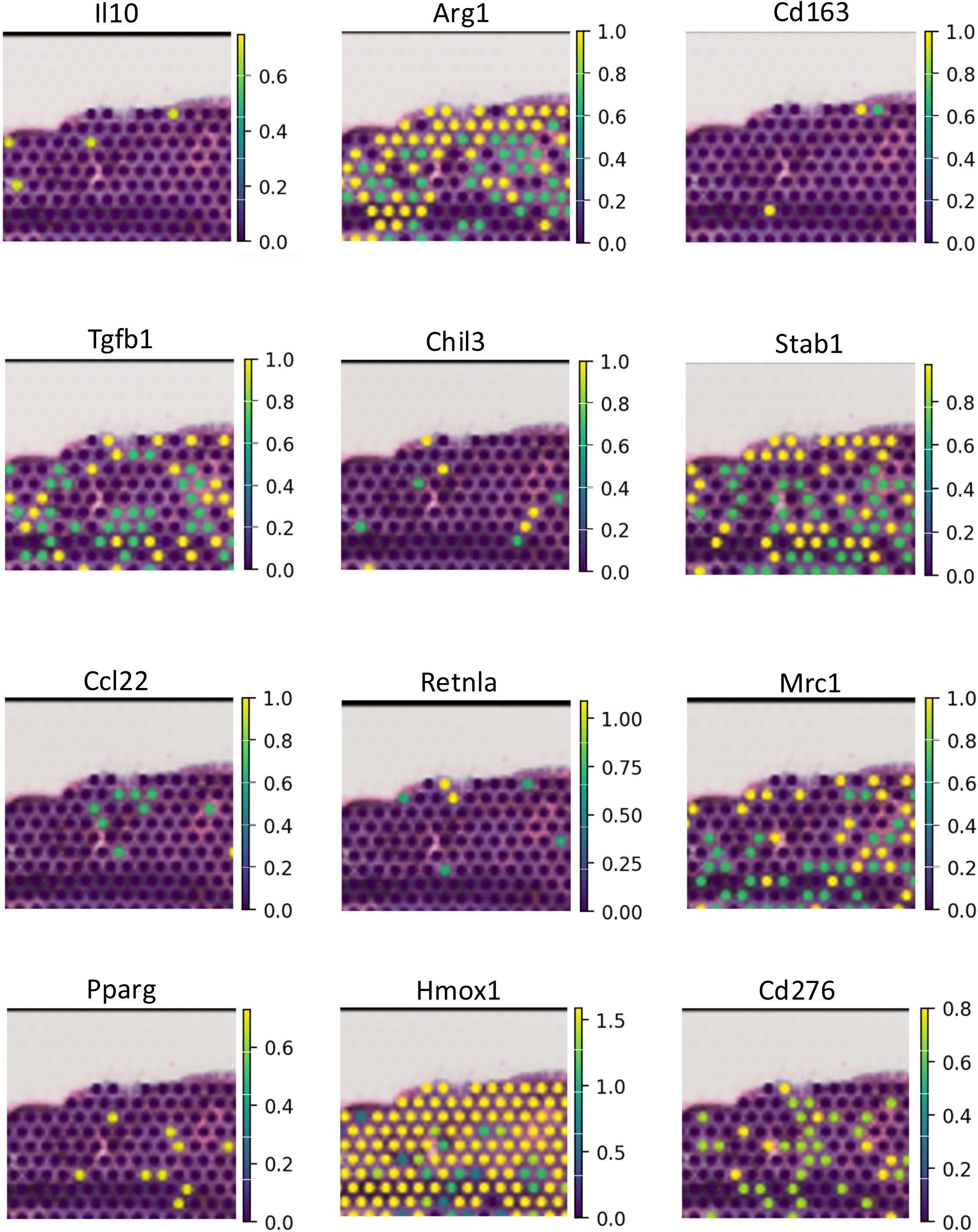
Spatial distribution of M2-associated markers in grid section A2. Expression patterns of 12 key M2 macrophage-associated markers in grid section A2 of the murine melanoma tumor. Each panel shows the spatial distribution of a different marker, with expression levels indicated by color intensity (yellow: high, green: intermediate, purple: low/none). Note the varying saturation scales (0-0.6, 0-0.8, 0-1.0, 0-1.5) across markers. Markers display distinct patterns ranging from high widespread expression (Hmox1, 0-1.5 scale) to strong localized expression (Arg1, Tgfb1, Stab1, Mrc1, 0-1.0 scale) to sparse or minimal expression (IL10, Cd163, Chil3, Ccl22, Retnla, 0-1.0 scale).

**Figure 5:**
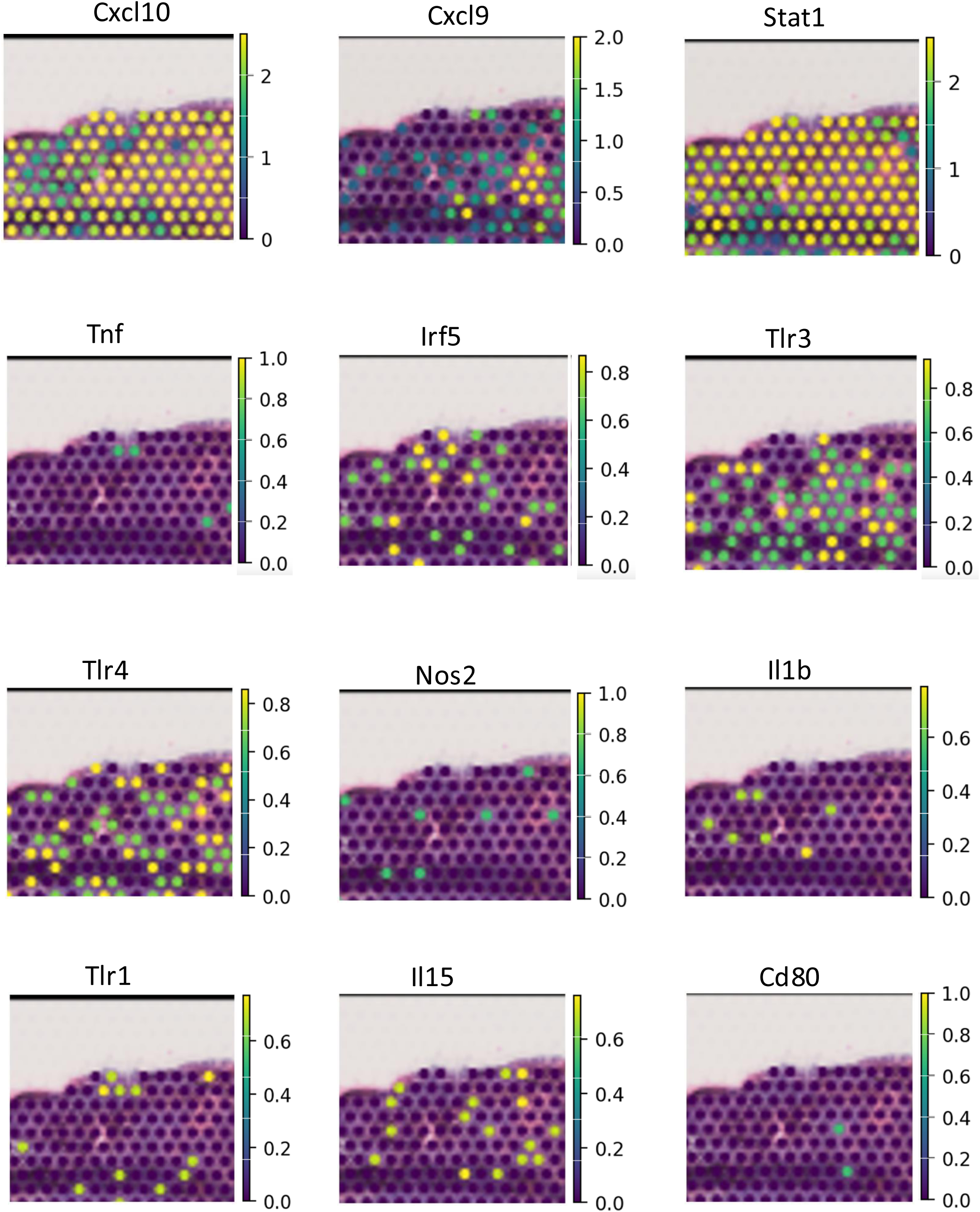
Spatial distribution of M1-associated markers in grid section A2. Expression patterns of 12 key M1 macrophage-associated markers in grid section A2 of the murine melanoma tumor. Each panel shows the spatial distribution of a different marker, with expression levels indicated by color intensity (yellow: high, green: intermediate, purple: low/none). Note the varying saturation scales across markers (0-2.0 for Cxcl10, Cxcl9, and Stat1; 0-0.8 for Irf5, Tlr3, and Tlr4; 0-1.0 for Tnf, Nos2, and Cd80; 0-0.6 for Tlr1, Il15, and Il1b). The patterns reveal striking heterogeneity in both expression levels and spatial organization. Highest expression is observed for Cxcl10 and Stat1 (0-2.0 scale), both showing abundant yellow spots throughout the section. In contrast, several key immune effectors including Tnf, Nos2, and Il1b show minimal expression despite using a more sensitive scale. Pattern recognition receptors (Tlr3, Tlr4) display moderate expression with notable spatial organization in the upper region, while Tlr1 shows minimal expression.

**Table 1:**
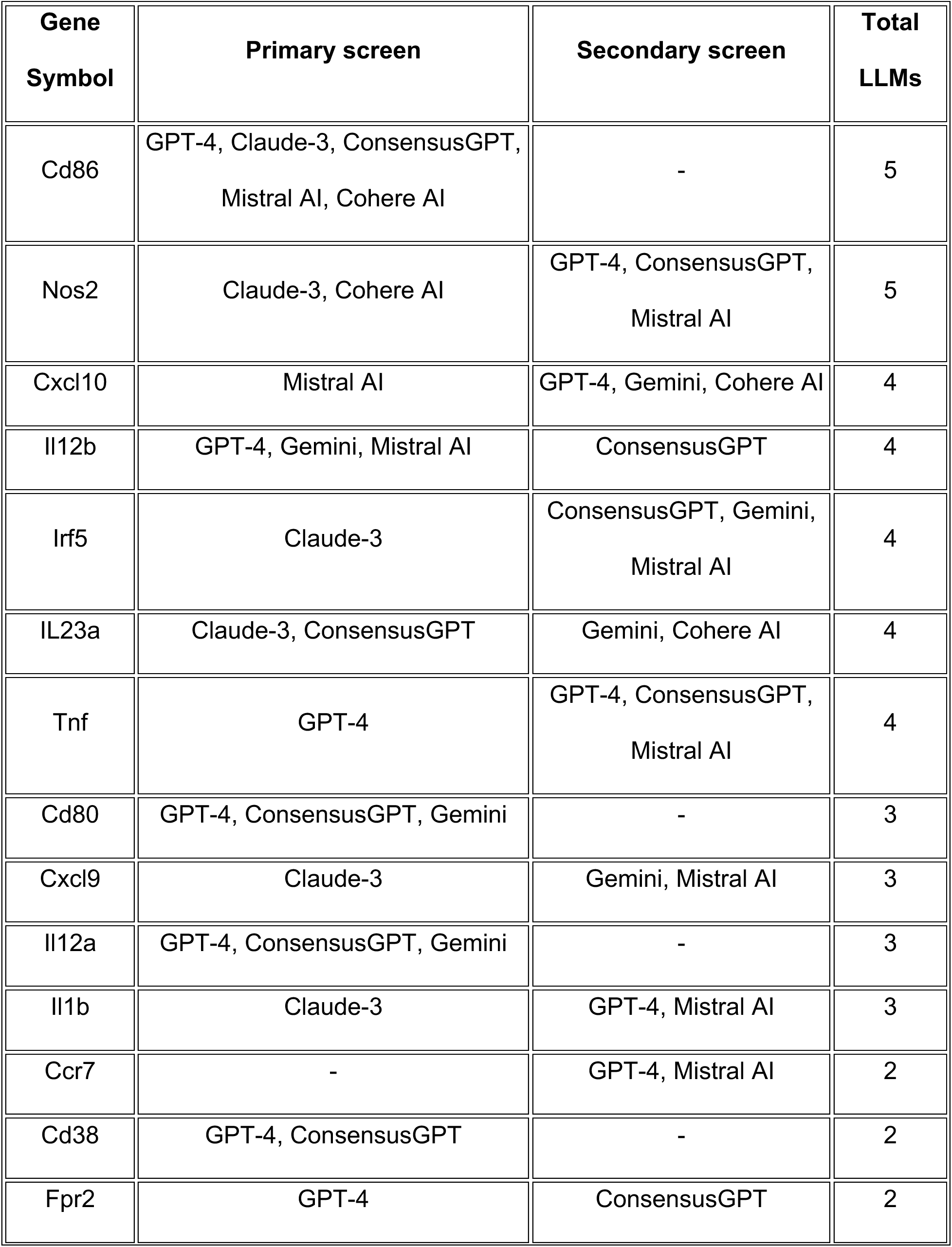
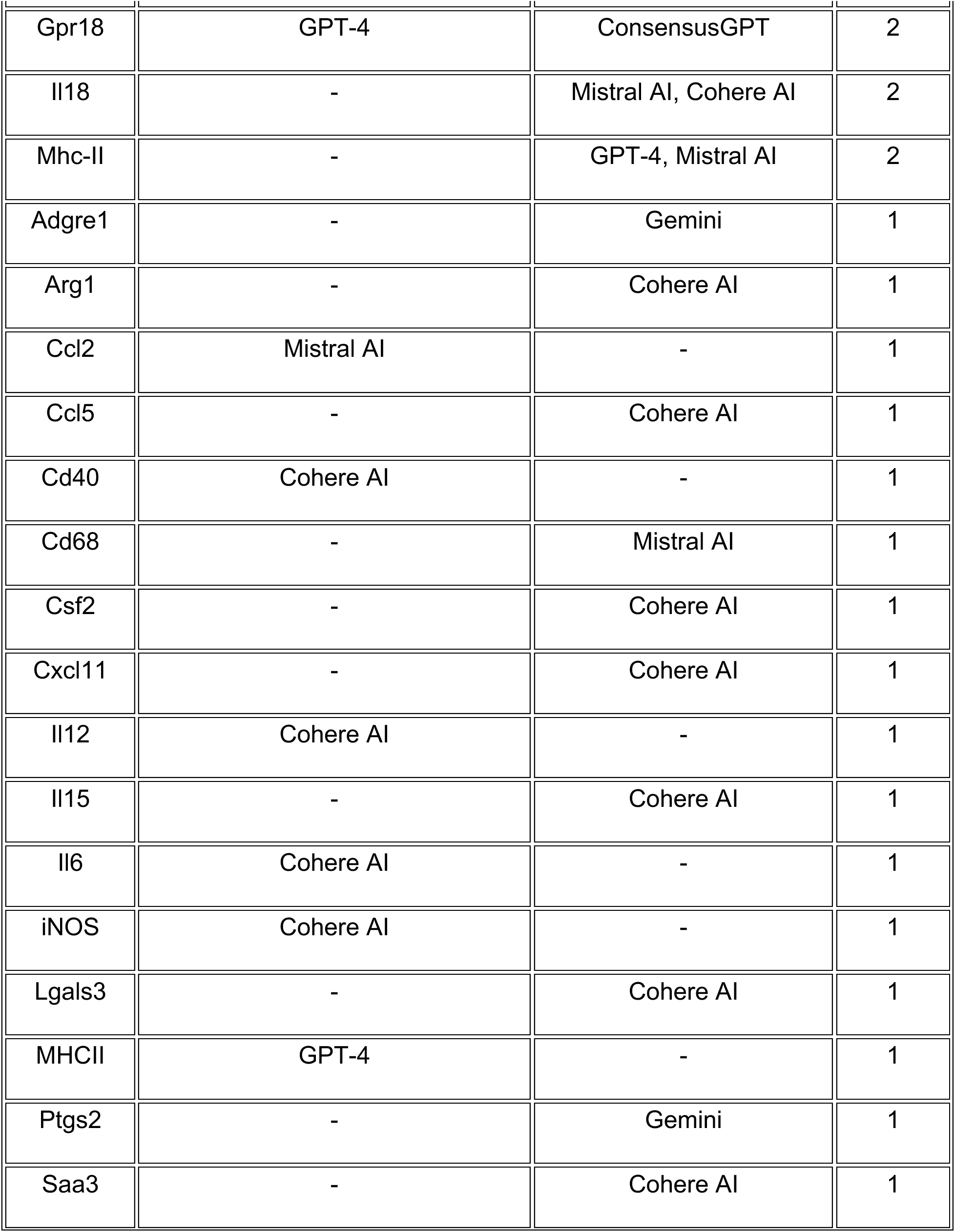
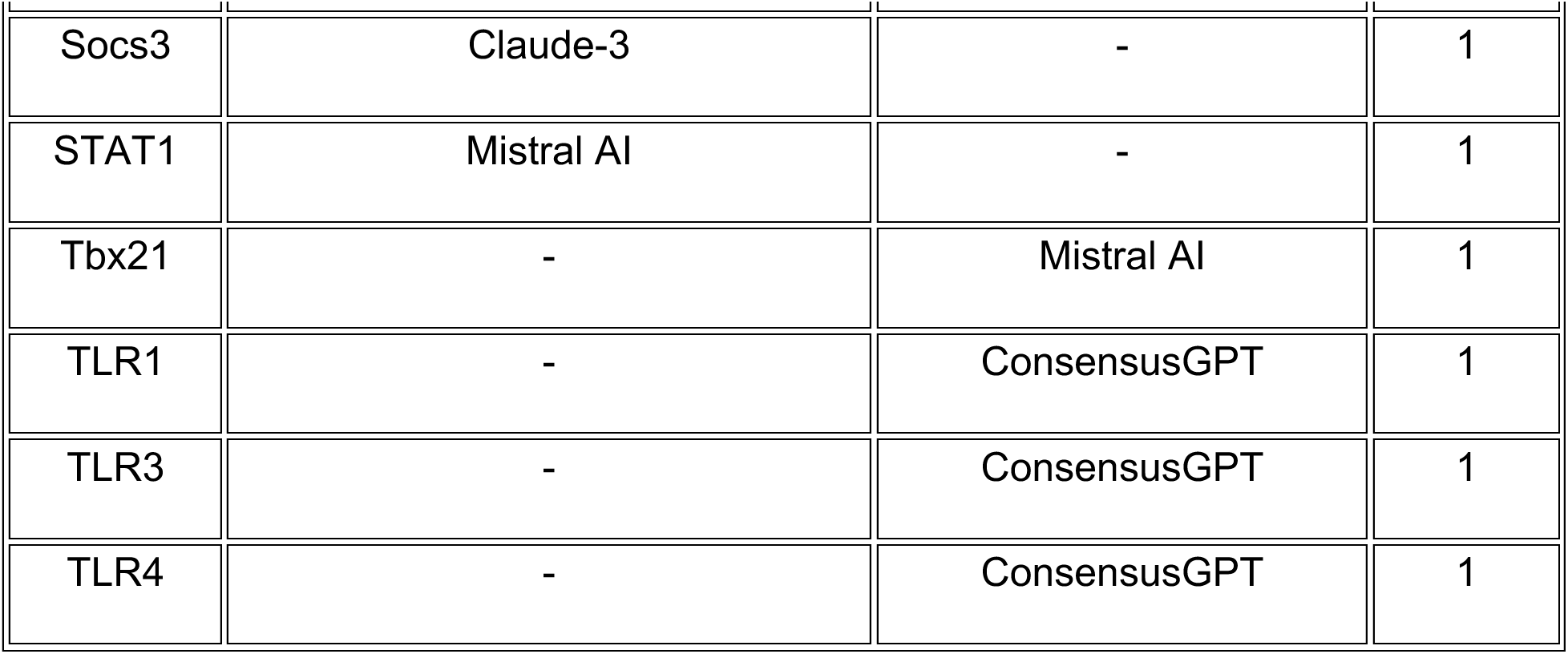
LLM Performance in Spatial Transcriptomics Spot Quantification.

| Gene Symbol | Primary screen | Secondary screen | Total LLMs |
|---|---|---|---|
| Cd86 | GPT-4, Claude-3, ConsensusGPT, Mistral AI, Cohere AI | - | 5 |
| Nos2 | Claude-3, Cohere AI | GPT-4, ConsensusGPT, Mistral AI | 5 |
| Cxcl10 | Mistral AI | GPT-4, Gemini, Cohere AI | 4 |
| Il12b | GPT-4, Gemini, Mistral AI | ConsensusGPT | 4 |
| Irf5 | Claude-3 | ConsensusGPT, Gemini, Mistral AI | 4 |
| IL23a | Claude-3, ConsensusGPT | Gemini, Cohere AI | 4 |
| Tnf | GPT-4 | GPT-4, ConsensusGPT, Mistral AI | 4 |
| Cd80 | GPT-4, ConsensusGPT, Gemini | - | 3 |
| Cxcl9 | Claude-3 | Gemini, Mistral AI | 3 |
| Il12a | GPT-4, ConsensusGPT, Gemini | - | 3 |
| Il1b | Claude-3 | GPT-4, Mistral AI | 3 |
| Ccr7 | - | GPT-4, Mistral AI | 2 |
| Cd38 | GPT-4, ConsensusGPT | - | 2 |
| Fpr2 | GPT-4 | ConsensusGPT | 2 |
| Gpr18 | GPT-4 | ConsensusGPT | 2 |
| Il18 | - | Mistral AI, Cohere AI | 2 |
| Mhc-II | - | GPT-4, Mistral AI | 2 |
| Adgre1 | - | Gemini | 1 |
| Arg1 | - | Cohere AI | 1 |
| Ccl2 | Mistral AI | - | 1 |
| Ccl5 | - | Cohere AI | 1 |
| Cd40 | Cohere AI | - | 1 |
| Cd68 | - | Mistral AI | 1 |
| Csf2 | - | Cohere AI | 1 |
| Cxcl11 | - | Cohere AI | 1 |
| Il12 | Cohere AI | - | 1 |
| Il15 | - | Cohere AI | 1 |
| Il6 | Cohere AI | - | 1 |
| iNOS | Cohere AI | - | 1 |
| Lgals3 | - | Cohere AI | 1 |
| MHCII | GPT-4 | - | 1 |
| Ptgs2 | - | Gemini | 1 |
| Saa3 | - | Cohere AI | 1 |
| Socs3 | Claude-3 | - | 1 |
| STAT1 | Mistral AI | - | 1 |
| Tbx21 | - | Mistral AI | 1 |
| TLR1 | - | ConsensusGPT | 1 |
| TLR3 | - | ConsensusGPT | 1 |
| TLR4 | - | ConsensusGPT | 1 |

**Table 2:** Top 15 M1 Macrophage-associated Markers.

| Rank | Gene Symbol | Average M1 Score | M1/M2 Ratio | Description |
|---|---|---|---|---|
| 1 | Il12 | 8.3 | 6.4 | Pro-inflammatory cytokine crucial for Th1 responses |
| 2 | Cxcl10 | 8.5 | 5.7 | Chemokine inducing T cell recruitment |
| 3 | Cxcl9 | 7.5 | 5.9 | IFN-γ induced chemokine promoting T cell trafficking |
| 4 | Stat1 | 8.3 | 4.9 | Transcription factor mediating IFN responses |
| 5 | Tnf | 8.0 | 4.2 | Pro-inflammatory cytokine with anti-tumor activity |
| 6 | Irf5 | 7.8 | 4.5 | Transcription factor promoting M1 polarization |
| 7 | Tbx21 | 6.5 | 7.0 | T-box transcription factor regulating Th1 development |
| 8 | Tlr3 | 6.8 | 5.6 | Pattern recognition receptor sensing viral RNA |
| 9 | Tlr4 | 7.3 | 4.6 | Pattern recognition receptor for bacterial LPS |
| 10 | Cxcl11 | 6.5 | 5.4 | Chemokine involved in T cell recruitment |
| 11 | Nos2 | 7.8 | 2.5 | Inducible nitric oxide synthase |
| 12 | Il1b | 7.3 | 3.0 | Pro-inflammatory cytokine |
| 13 | Tlr1 | 6.5 | 4.9 | Pattern recognition receptor |
| 14 | Il15 | 6.5 | 3.6 | Cytokine supporting NK and T cell responses |
| 15 | Cd80 | 6.8 | 1.9 | Co-stimulatory molecule |

**Table 3:**
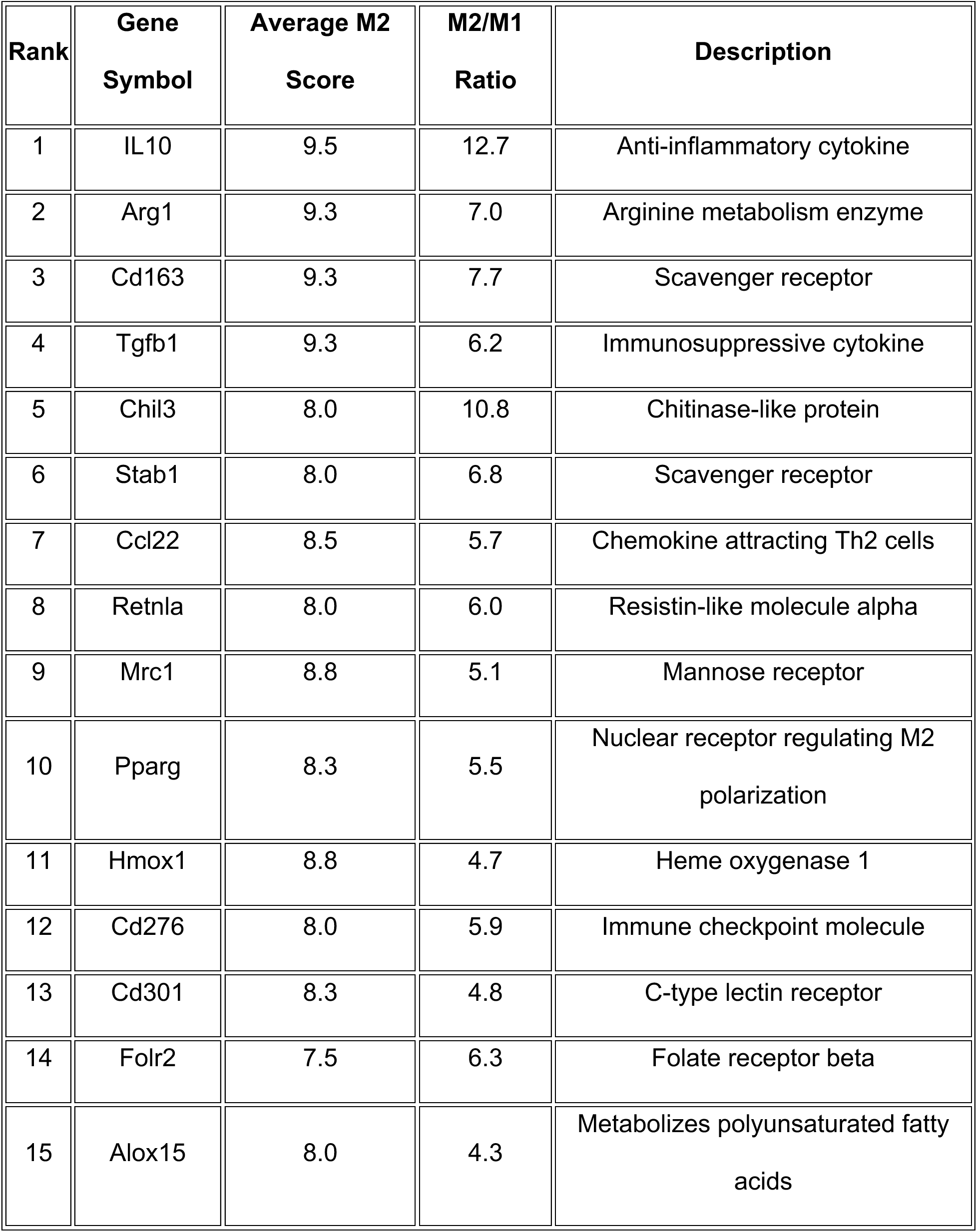
Top 14 M2 Macrophage-associated Markers.

| Rank | Gene Symbol | Average M2 Score | M2/M1 Ratio | Description |
|---|---|---|---|---|
| 1 | IL10 | 9.5 | 12.7 | Anti-inflammatory cytokine |
| 2 | Arg1 | 9.3 | 7.0 | Arginine metabolism enzyme |
| 3 | Cd163 | 9.3 | 7.7 | Scavenger receptor |
| 4 | Tgfb1 | 9.3 | 6.2 | Immunosuppressive cytokine |
| 5 | Chil3 | 8.0 | 10.8 | Chitinase-like protein |
| 6 | Stab1 | 8.0 | 6.8 | Scavenger receptor |
| 7 | Ccl22 | 8.5 | 5.7 | Chemokine attracting Th2 cells |
| 8 | Retnla | 8.0 | 6.0 | Resistin-like molecule alpha |
| 9 | Mrc1 | 8.8 | 5.1 | Mannose receptor |
| 10 | Pparg | 8.3 | 5.5 | Nuclear receptor regulating M2 polarization |
| 11 | Hmox1 | 8.8 | 4.7 | Heme oxygenase 1 |
| 12 | Cd276 | 8.0 | 5.9 | Immune checkpoint molecule |
| 13 | Cd301 | 8.3 | 4.8 | C-type lectin receptor |
| 14 | Folr2 | 7.5 | 6.3 | Folate receptor beta |
| 15 | Alox15 | 8.0 | 4.3 | Metabolizes polyunsaturated fatty acids |

**Table 4:** Spatial Analysis of M2 (Immunosuppressive) Macrophage Markers in Grid Section A2 of Murine Melanoma Tumor - Expression Patterns and Distribution.

| Marker | Total Spots | Yellow Spots | Green Spots | Purple Spots | Spatial Distribution | Marker Saturation Scale |
|---|---|---|---|---|---|---|
| IL10 | ~45 | 4 | 0 | ~41 | Yellow spots appear scattered near the upper edge of the section with no clear pattern. Purple spots dominate the majority of the area in a uniform distribution. No green spots observed. | 0.0-0.6 |
| Arg1 | ~45 | ~18 | ~12 | ~15 | Yellow spots concentrated in upper half of section forming clusters. Green spots distributed throughout, often adjacent to yellow spots. Purple spots more prevalent in lower portion. Clear gradient pattern from top to bottom. | 0.0-1.0 |
| Cd163 | ~45 | 3 | 1 | ~41 | Very sparse expression with only 3 yellow spots (2 clustered in upper right, 1 isolated in middle-left) and 1 green spot. Purple spots dominate throughout. | 0.0-1.0 |
| Tgfb1 | ~45 | ~15 | ~12 | ~18 | Yellow and green spots show relatively even distribution throughout the section, with slight clustering in upper portion. No strong gradient pattern. Purple spots interspersed throughout. | 0.0-1.0 |
| Chil3 | ~45 | 4 | 3 | ~38 | Very sparse expression pattern. Yellow spots scattered without clear pattern (1 upper left, 2 middle, 1 lower right). Few green spots dispersed across section. Purple spots dominant throughout. | 0.0-1.0 |
| Stab1 | ~45 | ~16 | ~14 | ~15 | Strong expression pattern with yellow spots concentrated in upper portion, particularly forming a distinct cluster. Green spots well distributed throughout. Clear spatial organization with higher expression in upper region. | 0.0-0.8 |
| Ccl22 | ~45 | 1 | ~8 | ~36 | Predominantly low expression with one yellow spot visible at the edge. Small cluster of green spots in upper middle region. Purple spots dominant throughout. | 0.0-1.0 |
| Retnla | ~45 | 2 | 4 | ~39 | Very sparse expression with two yellow spots in upper region and few scattered green spots. Purple spots dominant throughout. No clear spatial organization. | 0.00-1.00 |
| Mrc1 | ~45 | ~14 | ~12 | ~19 | Strong expression pattern with yellow and green spots well distributed throughout. Some clustering of yellow spots in upper portion. Similar pattern to Tgfb1 with good representation of both high and intermediate expression. | 0.0-1.0 |
| Pparg | ~45 | 8 | 0 | ~37 | Sparse expression with yellow spots scattered throughout, showing slight clustering in middle-lower region. No green spots observed. Purple spots dominant throughout. | 0.0-0.6 |
| Hmox1 | ~45 | ~30 | ~10 | ~5 | Highest overall expression among all markers. Yellow spots dominant throughout, particularly dense in upper portion. Some green spots interspersed. Very few purple spots. | 0.0-1.5 |
| Cd276 | ~45 | 6 | ~15 | ~24 | Moderate expression with more green than yellow spots. Green spots well distributed throughout the section. Yellow spots scattered with slight clustering in upper region. Purple spots still prominent. | 0.0-0.8 |
| Folr2 | ~45 | 4 | ~15 | ~26 | Moderate expression dominated by green spots. Few yellow spots scattered in upper region. Green spots well distributed throughout. Purple spots still make up majority. Similar pattern to Cd276 but with fewer yellow spots. | 0.0-0.8 |

**Table 5:** Spatial Analysis of M1 (Pro-inflammatory) Macrophage Markers in Grid Section A2 of Murine Melanoma Tumor - Expression Patterns and Distribution.

| **Marker** | **Total Spots** | **Yellow Spots** | **Green Spots** | **Purple Spots** | **Spatial Distribution** | **Marker Saturation Scale** |
|---|---|---|---|---|---|---|
| Cxcl10 | ~45 | ~25 | ~12 | ~8 | High overall expression with abundant yellow spots throughout, particularly concentrated in upper portion. Green spots well distributed throughout. Few purple spots. Shows a gradient with higher expression in upper region. | 0.0-2.0 |
| Cxcl9 | ~45 | ~8 | ~15 | ~22 | Moderate expression with mixed pattern. Yellow spots concentrated in lower right portion. Notable presence of blue spots (intermediate-low expression) throughout. Green spots well distributed. Clear spatial organization with higher expression in lower right region. | 0.0-2.0 |
| Stat1 | ~45 | ~30 | ~10 | ~5 | Very high overall expression with dominant yellow spots throughout. Green spots interspersed among yellow spots. Very few purple spots. More uniform distribution compared to Cxcl10 and Cxcl9, with strong expression across the entire section. | 0.0-2.0 |
| Tnf | ~45 | 0 | 4 | ~41 | Very sparse expression with only a few green spots scattered in the upper portion. No yellow spots present. Purple spots strongly dominant throughout the section. | 0.0-1.0 |
| Irf5 | ~45 | ~8 | ~12 | ~25 | Moderate expression with mix of yellow and green spots. Yellow spots showing some clustering in upper middle region. Green spots well distributed throughout. Purple spots still make up majority but less dominant than in Tnf. | 0.0-0.8 |
| Tlr3 | ~45 | ~12 | ~15 | ~18 | Moderate to high expression with well-balanced mix of yellow and green spots. Good distribution throughout the section with slight clustering in upper region. Purple spots less dominant than in previous markers. Even distribution between all three intensity levels. | 0.0-0.8 |
| Tlr4 | ~45 | ~10 | ~14 | ~21 | Moderate expression pattern similar to Tlr3 but slightly lower overall. Yellow and green spots well distributed throughout with some clustering in upper portion. Purple spots still prominent but not dominant. | 0.0-0.8 |
| Nos2 | ~45 | 0 | ~7 | ~38 | Very sparse expression with only a few green spots scattered throughout. No yellow spots present. Purple spots strongly dominant. Similar low expression pattern to Tnf. | 0.0-1.0 |
| Il1b | ~45 | 1 | ~5 | ~39 | Very sparse expression with one yellow spot and few green spots scattered throughout. Purple spots strongly dominant. Similar low expression pattern to Tnf and Nos2. | 0.0-0.6 |
| Tlr1 | ~45 | 2 | ~6 | ~37 | Sparse expression with two yellow spots and several green spots scattered throughout. No clear spatial organization. Purple spots strongly dominant. Similar pattern to Il1b but slightly more yellow spots. | 0.0-0.6 |
| Il15 | ~45 | ~10 | ~5 | ~30 | Low to moderate expression with more yellow than green spots. Yellow spots show some clustering in upper portion and edges. Purple spots still dominant but less so than other cytokines. | 0.0-0.6 |
| Cd80 | ~45 | 0 | 2 | ~43 | Very sparse expression with only two green spots visible. No yellow spots present. Purple spots overwhelmingly dominant. Lowest expression level observed among all M1 markers analyzed. | 0.0-1.0 |
| IL12 | ~45 | 0 | 0 | ~45 | No detectable expression throughout the section. | N/A |
| Tbx21 | ~45 | 0 | 0 | ~45 | No detectable expression throughout the section. | N/A |
| Cxcl11 | ~45 | 0 | 0 | ~45 | No detectable expression throughout the section. | N/A |

These results demonstrate Claude 3.5’s capability to systematically analyze complex spatial transcriptomics data, accurately quantifying not only expression levels but also capturing subtle spatial patterns and accounting for varying detection scales across markers.

#### Integration of Biological Knowledge in Spatial Transcriptomics Analysis

Having established Claude 3.5’s ability to quantify and map spatial expression patterns, we next investigated its capacity to integrate established immunological knowledge into its interpretation of the data. This would test whether an LLM could move beyond pattern recognition to provide mechanistically informed analysis of the tumor immune landscape. When prompted to factor in the functional significance of markers in its analysis, Claude 3.5 devised a knowledge-driven weighting approach, categorizing markers as highly significant (weight: 3), moderately significant (weight: 2), or less significant (weight: 1), with absent markers of high significance weighted accordingly (-3 for missing immunostimulatory markers, +3 for missing immunosuppressive markers). Using this framework, Claude 3.5 adjusted its initial immunological score for section A2 from -4 to - 5, supporting this revision with detailed analysis of several key biological factors:

1. The model identified the high expression of critical immunosuppressive regulators Arg1 and Tgfb1 as a primary factor, noting their synergistic effects in establishing a robust immunosuppressive environment through both direct signaling and metabolic regulation.
2. The absence of the key immune activator IL12 was highlighted as another critical factor, given its central role in Th1 responses and anti-tumor immunity.
3. Claude 3.5 also emphasized the metabolic impact of concurrent high Arg1 and Hmox1 expression, describing how their complementary effects on nutrient availability and metabolic products create a particularly hostile environment for effector T cells.
4. The model further demonstrated sophisticated pathway analysis proficiency by distinguishing between markers based on their position in signaling hierarchies. In the pro-inflammatory pathway, while the transcription factor Stat1 showed very high expression (30 yellow spots), its key downstream effectors exhibited strikingly low or absent expression: IL12 (undetectable), Tnf (4 green spots only), and IL1β (1 yellow spot), suggesting a "broken chain of command" in the immune-stimulatory response. Similarly, despite moderate expression of pattern recognition receptors Tlr3 and Tlr4 (12 and 10 yellow spots respectively), their downstream inflammatory mediators showed minimal activation. In contrast, the model identified complete and coherent activation of immunosuppressive pathways, exemplified by the coordinated expression from upstream regulators through effector molecules: high Tgfb1 expression (15 yellow spots) aligned with strong effector expression including Arg1 (18 yellow spots) and Hmox1 (30 yellow spots), while the M2-associated markers Mrc1, Stab1, and CD163 showed coordinated spatial distribution, particularly in the upper region. This pattern suggests not just immunosuppression, but a well-organized immunosuppressive state with disrupted pro-inflammatory signaling cascades.
5. Finally, Claude 3.5 appropriately weighted markers based on their direct functional impact, preventing over-interpretation of highly expressed but less functionally critical markers such as the M2-associated surface markers. The full chat transcript is available in Supplementary File 1.

#### Integrative Analysis Reveals Complex Organization of the Tumor Immune Microenvironment

Following quantitative and functional analyses, we challenged Claude 3.5’s ability to synthesize these findings into a comprehensive interpretation grounded in current immunological knowledge. When prompted to contextualize its observations and provide supporting evidence from literature, the model generated the following assessment: "The immunological landscape of section A2 reveals a sophisticated organization of immune evasion that both confirms and extends our current understanding of the tumor microenvironment. At its core, we observe the established paradigm of M2-polarized tumor-associated macrophages (10), but with important nuances in spatial organization and signaling hierarchies. The dominant feature is a coordinated metabolic barrier, primarily through Arg1 and Hmox1 co-expression. While these enzymes’ individual roles in immune suppression are well-documented (11,12), their spatial co-organization suggests a more sophisticated suppressive mechanism than previously appreciated.

Perhaps most intriguing is the preservation of upstream M1 signaling (high Stat1) despite compromised effector functions, reminiscent of the "stunned" immune phenotype described in chronic infections (13). This diverges from the simple M1/M2 dichotomy traditionally described in tumor immunology (14). The spatial organization of immunosuppressive mechanisms aligns with emerging concepts of "immune niches" in tumors (15), though our observations suggest more complex organizational principles than previously recognized. These findings have therapeutic implications, supporting the growing consensus that effective immunotherapy may require targeting multiple aspects of the tumor immune environment (16), with particular attention to metabolic reprogramming and signal cascade restoration."

This analysis demonstrates Claude 3.5’s ability not only to process complex spatial molecular data but also to integrate this information with established biological knowledge, generate novel insights, and support its interpretations with relevant literature. This capability suggests significant potential for LLMs in advancing our understanding of complex tissue organization and function.

## DISCUSSION

In this study, we explored the potential of Large Language Models to assist in the interpretation of spatial transcriptomics data, focusing specifically on understanding the complex immune landscape of the tumor microenvironment. Our systematic evaluation began with benchmarking multiple LLMs for their ability to analyze spatial gene expression patterns, revealing that only Claude 3.5 Sonnet demonstrated sufficient capability for this complex task. We then developed a structured analytical workflow leveraging Claude 3.5 to interpret the spatial organization of immune responses in a murine melanoma model. Starting with the identification and prioritization of M1 and M2 macrophage markers through knowledge-driven scoring, we proceeded to analyze their spatial distribution patterns in defined tumor regions. The model showed remarkable ability not only to quantify expression patterns but also to integrate this information with established immunological knowledge, ultimately providing sophisticated interpretation of the tumor immune landscape. This was exemplified in the analysis of a given grid section (A2), where Claude 3.5 identified a well-organized immunosuppressive state characterized by coordinated metabolic barriers and disrupted pro-inflammatory signaling cascades, findings that both aligned with and extended current understanding of tumor immunology.

The high level of proficiency demonstrated by Claude 3.5 in interpreting spatial transcriptomics data was unexpected, particularly its ability to accurately quantify spot numbers while accounting for varying saturation scales and to characterize complex spatial distributions. However, this capability represents a frontier skill in LLM development, as evidenced by our benchmarking results where only two of six tested models could take on this task, with varying degree of success. This finding aligns with recent studies exploring LLMs’ ability to interpret scientific visualizations. For instance, Royer et al. (9) highlighted the emerging capabilities of LLMs in bioimage analysis while noting significant variation in performance across models and emphasizing the need for careful evaluation of these systems when applied to scientific image interpretation tasks. The ability to accurately interpret spatial molecular data thus appears to be a sophisticated capability limited to the most advanced models, likely resulting from recent improvements in multimodal training approaches and visual-spatial reasoning. Notably, we observed rapid progress in this domain during our study period, with newer model iterations showing marked improvements in spatial pattern recognition accuracy. For example, the late October 2024 update of Claude 3.5 demonstrated enhanced performance in spot quantification compared to its previous version. And one of the most improved models during this period being Google Gemini. This trajectory of quick improvement suggests significant potential for broader application of LLMs in analyzing complex biological data visualizations in the near future.

Our approach to marker prioritization and immune status scoring builds upon scoring frameworks we previously developed for systematic evaluation of biomarkers and interpretation of high-dimensional molecular data (8,17). While these earlier studies demonstrated the utility of LLMs in knowledge-driven candidate gene selection, the current work extends this capability to assisting with the interpretation of complex spatial data. Particularly noteworthy was Claude 3.5’s ability to not only quantify expression patterns but to integrate this information with extensive knowledge about marker function and hierarchy in the tumor microenvironment. This was especially evident in the model’s transition from initial expression-based scoring to functionally-weighted analysis of section A2, where it demonstrated sophisticated understanding of both molecular mechanisms and spatial organization. For instance, the model independently devised a weighting system that appropriately prioritized master regulators of immune function while accounting for pathway relationships and metabolic effects. This capacity for knowledge integration was further demonstrated by its ability to identify significant patterns such as the spatial co-organization of metabolic regulators Arg1 and Hmox1, and to contextualize these findings within current understanding of tumor immunology (14). Such ability to synthesize spatial molecular data with broad biological knowledge represents a significant advance in the application of LLMs to biomedical data interpretation, suggesting potential applications beyond spatial transcriptomics to other complex molecular profiling approaches. Most notably, the model demonstrated emergent analytical capabilities by developing novel approaches to data interpretation without explicit prompting. A striking example was its analysis of signaling pathway "completeness," where it contrasted the disrupted pro-inflammatory cascade (high Stat1 expression despite low downstream effector activity) with fully operational immunosuppressive pathways. This type of analysis arose spontaneously from the model’s integration of expression data with pathway knowledge, offering interesting perspectives on the organization of the tumor immune microenvironment. Such emergent analytical capabilities suggest that LLMs might not only assist in data interpretation but also contribute novel insights for understanding complex biological systems.

Several limitations of our study should be noted. While Claude 3.5 demonstrated impressive capability in analyzing spatial transcriptomics data, specialized image analysis tools would likely provide superior accuracy and throughput for basic tasks such as spot counting and intensity measurements. However, the unique value of LLMs lies in their ability to combine competent visual analysis with extensive biomedical knowledge integration - a capability that mirrors human expert analysis but operates at much greater speed and scale. This combination permits rapid, context-aware interpretation of spatial molecular data that goes beyond simple quantification to provide mechanistic insights and biological interpretation. Furthermore, It is important to emphasize that this study represents a proof of concept, establishing a blueprint for LLM-assisted spatial transcriptomics analysis. The current workflow, which relied on a chat interface for development and testing, is not readily scalable for high-throughput analysis. Future implementation will require development of specialized applications that interface with foundational models through APIs, allowing for automated processing of multiple tissue sections and integration with existing analytical pipelines. Such development should also address challenges of reproducibility and standardization across different LLM versions and implementations.

In conclusion, the emergence of spatial transcriptomics technologies is transforming our understanding of tissue organization and function, but the complexity of the data generated poses significant analytical challenges. Our proof of concept demonstrates that advanced LLMs can assist in the interpretation of these complex datasets by combining pattern recognition capabilities with biological knowledge integration. This capability is particularly relevant given the increasing adoption of spatial transcriptomics approaches across biomedical research. While significant development work remains to be done, the analytical framework we established could help accelerate the translation of spatial transcriptomics data into biological insights.

## ACKNOWLEDGEMENTS

This work was supported by the Jackson Laboratory’s CATch program and The Jackson Laboratory’s Data Science Initiative. The V Foundation (V2021-036) and the AAI Careers in Immunology Fellowship Program provided support to CC. The authors acknowledge support from the following JAX Scientific Services: Histology Service, Single-Cell Biology, Genome Technology, and Flow Cytometry, which are partially supported by The JAX Cancer Center Support Grant (P30CA034196) and the Computational Sciences Department.

